# Kinesin-1 promotes centrosome clustering and nuclear migration in the *Drosophila* oocyte

**DOI:** 10.1101/2022.02.16.480671

**Authors:** Maëlys Loh, Fred Bernard, Antoine Guichet

## Abstract

Accurate positioning of the nucleus is essential. Microtubules and their associated motors are important players in this process. Although nuclear migration in *Drosophila* oocytes is controlled by microtubule, a role for microtubule-associated molecular motors in nuclear positioning has yet to be reported. We characterize novel landmarks that allow a precise description of the pre-migratory stages. Using these newly defined stages, we report that, prior to migration, the nucleus moves from the oocyte anterior side toward the center and concomitantly the centrosomes cluster at the posterior of the nucleus. In absence of Kinesin-1, centrosome clustering is impaired and the nucleus fails to position and migrate properly. The maintenance of a high level of Polo-kinase at centrosomes prevents centrosome clustering and impairs nuclear positioning. In absence of Kinesin-1, SPD2 an essential component of the pericentriolar material is increased at the centrosomes, suggesting that Kinesin-1 associated defects result from a failure to reduce centrosome activity. Consistently, depleting centrosomes rescues the nuclear migration defects induced by Kinesin-1 inactivation. Our results suggest that Kinesin-1 controls nuclear migration in the oocyte by modulating centrosome activity.

**Summary statement:** In this study, we identified a crucial role of Kinesin-1 in centrosome clustering required for nuclear positioning and migration in the *Drosophila* oocyte.

## Introduction

The cytoskeleton plays a central role in nuclear positioning, which regulates many cellular and developmental systems. Microtubules (MTs) as well as associated motor proteins are instrumental in the underlying mechanisms of this positioning. In many cases, MTs participate to localize the nucleus in close association with a centrosome, acting as MT organizing center. The MT plus-end directed motor, Kinesin-1, plays an essential function in the nuclear positioning in several cellular contexts (Duncan and Warrior, 2002; Folker et al., 2013; Fridolfsson and Starr, 2010; Januschke et al., 2002; Metzger et al., 2012; Meyerzon et al., 2009; Roux et al., 2009; Splinter et al., 2010; Williams et al., 2014; Wilson and Holzbaur, 2012; Wilson and Holzbaur, 2015). Kinesin-1 is a hetero-tetramer, composed of a dimerized Kinesin heavy chains (Khc) and two regulatory Kinesin light chains (Klc) (Verhey et al., 2011). Kinesin-1 can affect nuclear positioning via different mechanisms. For example, Kinesin-1 can transport the nucleus as cargo and drive its displacement along MTs towards their plus ends via an interaction between Klc and nuclear envelope (NE) associated proteins such as Nesprins (Meyerzon et al., 2009; Roux et al., 2009). Alternatively, Khc can control nuclear positioning independently of Klc by crosslinking MTs via a C-terminal MT binding site or indirectly through an association with the MT associated protein (MAP) Ensconsin/Map7 (Metzger et al., 2012). In the *Drosophila* oocyte, asymmetric positioning of the nucleus is crucial for the organization of MT-based transport which controls, among other things, the asymmetric localization of mRNAs that encode determinants of the polarity axes of the future embryo (González-Reyes et al., 1995; Guichet et al., 2001; Januschke et al., 2006; Roth et al., 1995; Swan et al., 1999).

During the 14-stage process of oogenesis, the oocyte is specified by a group of 16 interconnected germ cells, while the remaining 15 cells differentiate into nurse cells (NCs) (Huynh and St Johnston, 2004). This germline cyst is surrounded by epithelial follicle cells that form the egg chamber. The oocyte is positioned at the posterior of the egg chamber in contact with the follicle cells. (Fig; 1B) (Huynh and St Johnston, 2004). The oocyte contains at least 16 centrosomes, as in early oogenesis the centrosomes of the 15 NCs migrate through the ring canals into the oocyte to eventually form a cluster located between the nucleus and the posterior plasma membrane of the oocyte (Bolívar et al., 2001; Mahowald and Strassheim, 1970; Nashchekin et al., 2021; Pimenta-Marques et al., 2016). During mid-oogenesis, this cluster co-migrates with the nucleus and remains asymmetrically localized in close vicinity to the nucleus (Januschke et al., 2006; Tissot et al., 2017; Zhao et al., 2012). Then, the centrosomes gradually lose their pericentriolar materials (PCM), leading to their elimination during late oogenesis (Pimenta-Marques et al., 2016).

Pioneering studies have revealed that the movement of the *Drosophila* oocyte nucleus is mediated by MTs (Koch and Spitzer, 1983). More recent work further showed that a dual relationship exists between the nucleus and the MTs within the oocyte. Nuclear positioning influences the MT network organization in the oocyte. Additionally, the MTs are instrumental in the nuclear migration (Januschke et al., 2006; Tissot et al., 2017; Zhao et al., 2012). After receiving an unknown signal from the posterior follicular cells (González-Reyes et al., 1995; Roth et al., 1995), between stages 6 and 7, the nucleus of the oocyte migrates from the center to the anterior side of the oocyte and is subsequently maintained at the boundary between the plasma membrane of the anterior margin and the lateral membrane, for review see (Bernard et al., 2018). Migration of the nucleus is achieved by MTs pushing forces exerted on the NE (Tissot et al., 2017; Zhao et al., 2012). We have further reported that distinct molecular cues act in complementary fashion during this process (Tissot et al., 2017). One is associated with the MAP Mushroom-body defect (Mud), which is asymmetrically located at the NE of oocyte nucleus. The second cue corresponds to the centrosomes.

However, the mechanisms ensuring the onset of nuclear migration remain unknown. Furthermore, although MT involvement has been clearly demonstrated, the potential involvement of MT-associated motors, particularly Kinesin-1 have not been identified in the oocyte nuclear migration. Kinesin-1 has been reported to be necessary only for the positioning maintenance of the nucleus after its asymmetric migration. Moreover, only Khc but not Klc is required for this process (Duncan and Warrior, 2002; Januschke et al., 2002; Loiseau et al., 2010; Palacios and St Johnston, 2002; Williams et al., 2014).

In this study, by detailing the developmental stages preceding the migration of the nucleus in the *Drosophila* oocyte, we have further characterized the mechanisms controlling the onset of nuclear migration and reveal a role for Kinesin-1. We found, despite differences in function at later stages, that both subunits of Kinesin-1, Khc and Klc, are involved for nuclear positioning and migration. We further show that Kinesin-1 is required for centrosome clustering at the onset of the nuclear migration and that the two processes are correlated. We found that Kinesin-1 modulates the level of PCM at the centrosomes and propose that the motor controls oocyte nuclear positioning and migration by down-regulating centrosomal activity.

## Results

### Analysis of nuclear positioning prior to migration

The nature of mechanisms ensuring nuclear migration onset remains unknown, and more generally nucleus position prior migration has been poorly studied and commonly described as roughly posterior within the oocyte. Therefore, we sought to better characterize nucleus position and the development of the oocyte and egg chamber from stages 5 to 7, which precede and overlap with nuclear migration, respectively. Recently, there was a reported a method to assign egg-chamber stages based on NC diameter (Chen et al., 2019). We applied this methodology to two of the four NCs in contact with the oocyte in order to describe the period of nuclear pre-migration (Fig. 1A, B). We found that at stage 5 the nucleus is always anteriorly positioned in the oocyte, in contact with or in close vicinity of the anterior membrane of the oocyte, after which the nucleus will progressively become more centered (see methods). Indeed, even if oocyte development is a continuous process and our measured NC diameters reflects this continuity, when we investigated at the nucleus positioning, we found that there are distinct populations of stage 6 oocytes, the early stage 6 associated with smaller NC diameters and the late-stage 6 associated with larger NC diameters, which we decided to name 6A and 6B, respectively. Thus, the measurement of the NC diameter (Fig. 1 C, D) together with the egg chamber aspect ratio and the oocyte shape (see methods) allowed us to distinguish four steps, refining this developmental phase through the stages 5, 6A, 6B and 7. At stage 6A, most of oocytes exhibit an anteriorly positioned nucleus in the oocyte, even if there are some instances of central positioning. At stage 6B, the nucleus is mostly found in a central position and at stage 7 the migration is complete, and the nucleus is in contact with both the anterior and the lateral plasma membranes of the oocyte (Fig. 1 E, F).

**Figure 1.**
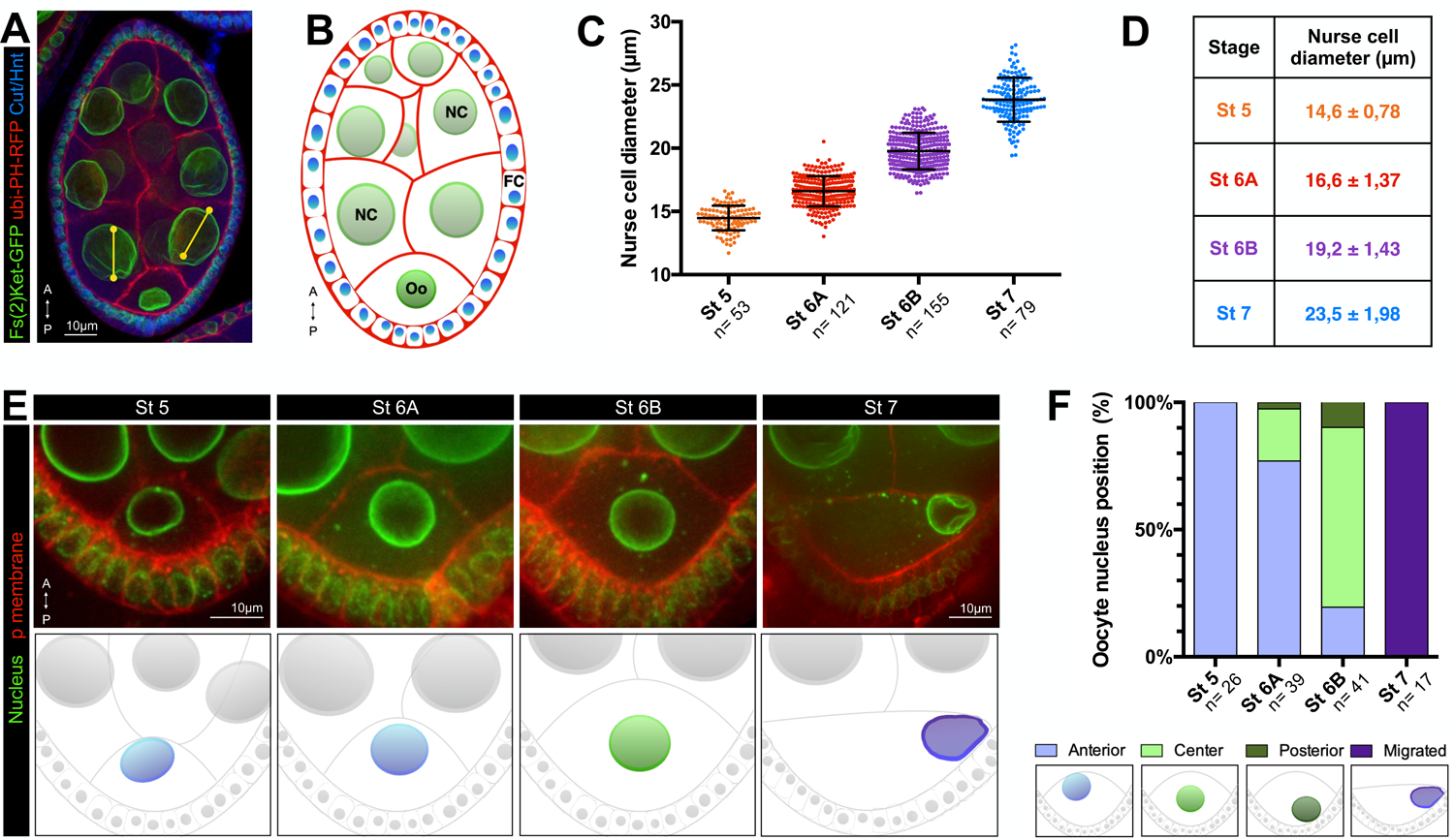
Oocyte staging and characterization of the nucleus positioning prior to migration. **(A-B)** Stage 6 egg chamber oriented with anterior (A) at the top and posterior (P) at the bottom, expressing *Fs(2)Ket-GFP* to label nuclei (green) and *pubi-PH^PLC∂1^-RFP* to label plasma membrane (red), stained with Cut antibody (blue) (A) and schematic diagram highlighting the nuclei of the different cell types: the follicular cells (FC) in blue, the nurse cells (NC) in light green and the oocyte (Oo) in dark green (B). Scale bar: 10µm. **(C)** Distribution of NC diameters shows a progressive increase in size and allows the categorization of 4 different stages (n indicates the number of analyzed egg chambers but dots correspond to each measured nuclei). Means ± s.e.m for each stage are indicated in D. **(E)** Stage 5 to 7 egg chambers, expressing *Fs(2)Ket-GFP* to label nuclei (green) stained with Cellmask to reveal plasma membranes (red). Representative examples of the different nuclear positions at stages 5, 6A, 6B and 7. The oocytes are oriented with anterior (A) at the top and posterior (P) at the bottom. Scale bar: 10µm. Schematic representation of the image above, with a color code illustrating the oocyte nuclear position, respectively pale blue for an anterior, pale green for center and violet for migrated to the intersection between anterior plasma membrane and the lateral plasma membrane. **(F)** Distribution of nucleus positions at the different stages. Positions have been categorized and color-coded as examplified in E. n indicates the number of analyzed egg chambers. See Sup Table 1 for detailed values of the quantifications.

In order to confirm the distinction between those stages, we decided to investigate a criterion independent of the germline and thus took advantage of the cell cycle switch that occurs in the surrounding follicular epithelium (Rowe et al., 2020) simultaneously to the nuclear migration in the oocyte. Cut and Hindsight (Hnt) are two transcription factors typically used as specific markers of the mitotic and endocycle cycles, respectively (Sun and Deng, 2005; Sun and Deng, 2007). Accordingly, before oocyte nuclear migration (stage 6), the follicular cells express Cut, and after completion of nuclear migration (stage 7) they express Hnt (Sup Fig 1). However, within stage 6 follicles, there are intermediate cases where Cut and Hnt are not expressed, further supporting two distinct phases within stage 6.

### Khc and Klc subunits are differentially required for nuclear positioning

Using these stage definitions, we looked for factors that are required for initiating nuclear migration. We found that RNAi mediated inactivation of *Khc* in the oocyte impairs the position of the nucleus. In this context, the nucleus remains anteriorly positioned at stages 6A and 6B and subsequently fails to migrate at stage 7 (Fig. 2A-B). Similar results were obtained with a second RNAi line directed against *Khc* (Fig. 2C). We also confirm our results by generating germline mitotic clones. Induction of GFP/FRT clones of the *Khc^27^* mutant allele, a *Khc* null allele (Januschke et al., 2002), revealed similar phenotypes to the RNAi knockdown conditions, thus confirming that Khc is necessary for nuclear migration and that its loss results in non-centered nucleus at stage 6B (Fig. 2 D, E).

**Figure 2.**
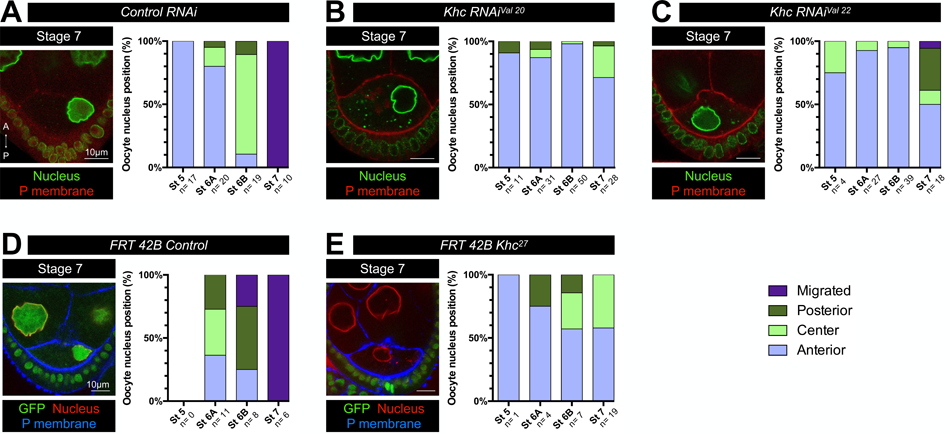
Khc is required for nucleus positioning and migration. Representative image of stage 7 egg-chambers and distribution of nucleus positions at the different stages. Positions have been categorized and color-coded as anterior in pale blue, center in pale green, posterior in dark green and migrated in purple. n indicates the number of analyzed egg chambers. **(A-C)** RNAi mediated analysis of nucleus positions where control RNAi (*UASp-CG12699-RNAi – See Methods*) (A) *Khc-RNAi^Val20^* (B) and *Khc-RNAi^Val22^*(C) have been expressed using the *mat-αtub-Gal4* driver combined with *Fs(2)Ket-GFP* to label nuclei (green) and *pubi-PH^PLC∂1^-RFP* to label plasma membrane (red). **(D-E)** GFP/FRT clonal analysis of nucleus positions in control egg chambers (*Khc^27^* heterozygous) (D) and *Khc^27^* mutant egg chambers revealed by the absence of GFP in germline nuclei (E). Nuclei and plasma membranes are revealed by WGA and SiR-actin staining, respectively. See Sup Table 1 for detailed values of the quantifications.

Subsequently, we decided to test the role of Klc, the non-motor subunit of Kinesin-1 in nuclear positioning. Similarly, to *Khc* down-regulation, RNAi mediated knockdown of *Klc* in the oocyte leads to mispositioned nuclei that remain at the anterior of the oocyte at stages 6A and 6B. In this genetic background, stage 7 egg chambers display non-migrated nuclei (Fig.3 A-C). This indicates that Klc is also required for the nucleus positioning. Furthermore, a GFP/FRT-mediated clonal analysis with two distinct *Klc* alleles confirmed these results (Fig.3 D-F). However, in mutant contexts for *Klc*, we noted that the nuclei eventually migrate, as we can detect some correctly positioned nuclei at stage 9 and beyond (Sup. Fig. 2 C-E), consistent with previously published studies (Palacios and St Johnston, 2002). The latter result contrasts with *Khc* mutant oocytes that still display nuclear positioning defects at stage 9 and beyond (Sup. Fig. 2 A, B). This difference underlines different requirements for both subunits, consistently with previous reports and hypotheses arguing for Kinesin-1 functions independent of Klc (Loiseau et al., 2010; Palacios and St Johnston, 2002), but could also reflect different subcellular distributions of Khc and Klc. Indeed, when we assessed their respective intracellular location using GFP-fusion proteins, we found that both subunits are evenly distributed in the oocyte cytoplasm but Khc is additionally enriched around the nucleus and at the centrosomes, while Klc is not (Sup Fig. 3 A, B, C).

**Figure 3.**
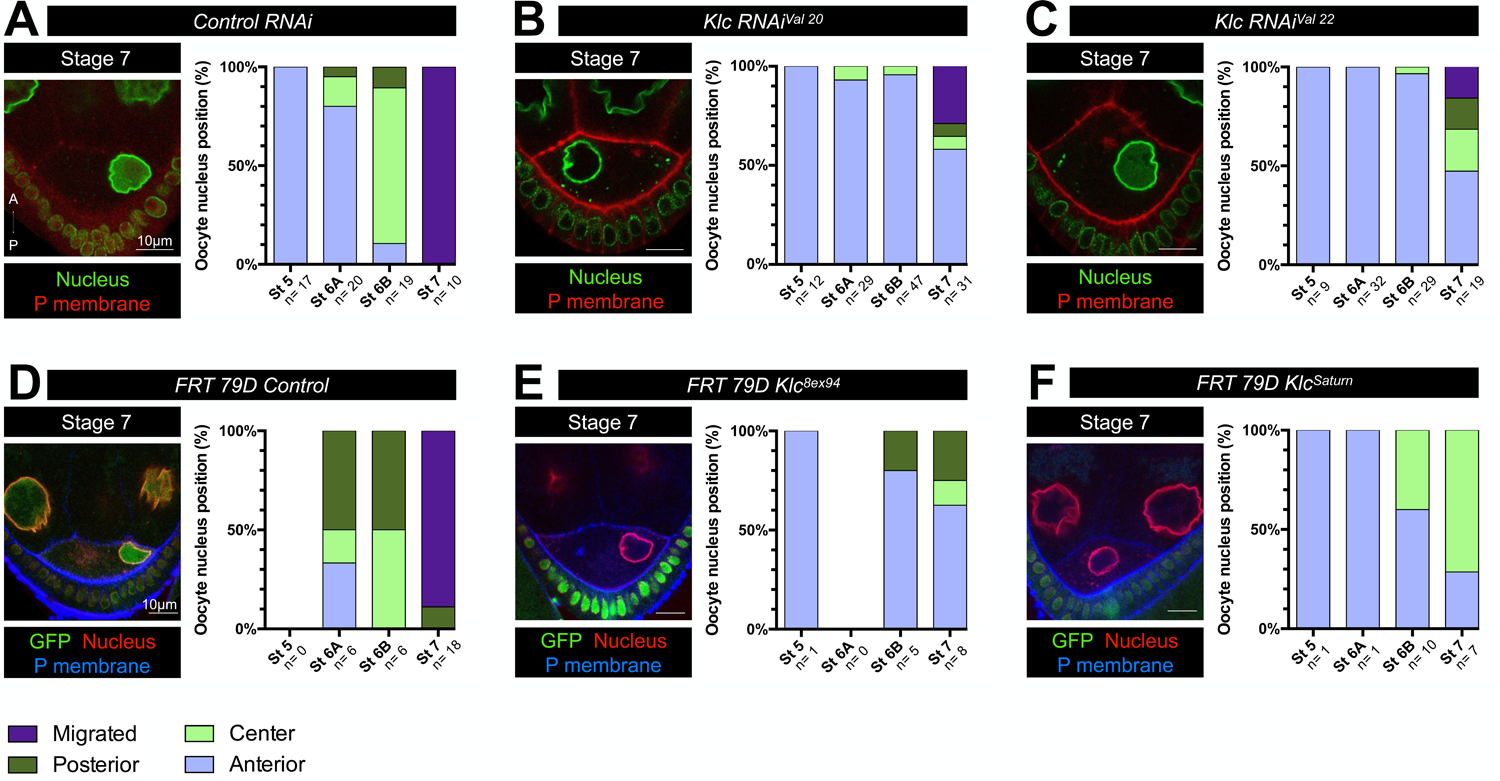
Klc is required for the nucleus positioning and efficient migration. Representative image of stage 7 egg-chambers and distribution of nucleus positions at the different stages. Positions have been categorized and color-coded as anterior in pale blue, center in pale green, posterior in dark green and migrated in purple. n indicates the number of analyzed egg chambers. **(A-C)** RNAi mediated analysis of nucleus positions where control RNAi (*UASp-CG12699-RNAi – See Methods,* identical panel to Figure 2A) (A) *Klc-RNAi^Val20^* (B) and *Klc-RNAi^Val22^* (C) have been expressed using the *p_mat_αtub-Gal4* driver combined with *Fs(2)Ket-GFP* to label nuclei (green) and *pubi-PH^PLC∂1^-RFP* to label plasma membrane (red). **(D-F)** GFP/FRT clonal analysis of nucleus positions in control egg chambers (*Klc* heterozygous) (D), *Klc^8ex94^*(E) and *Klc^Saturn^* (F) mutant egg chambers revealed by the absence of GFP in germline nuclei. Nuclei and plasma membranes are revealed by WGA and SiR-actin staining, respectively. See Sup Table 1 for detailed values of the quantifications. The data shown in Fig. 2A and Fig. 3A are identical.

Altogether, these results show that both Kinesin-1 subunits are required for early positioning and migration onset between stages 6 and 7, but also that a Khc-dependent process, although less efficient, is sufficient to ensure the nuclear migration and lead to delayed asymmetric positioning.

### Khc and Klc subunits differentially affect the MTs

Kinesin-1 has been shown to interfere directly with MT organization (Daire et al., 2009; Drechsler et al., 2020; Nieuwburg et al., 2017), especially through its ability to crosslink MTs (Lu et al., 2016; Metzger et al., 2012). Hence, we wondered if the nuclear migration defects observed in absence of Khc and Klc could be explained by defects in MT organization. We therefore decided to quantify MT density from stage 5 to 7, in *Khc* and *Klc* knockdown contexts. To do so, we assessed MT organization by quantifying the signal of Jupiter-GFP, a MAP that binds along the length of MTs (Baffet et al., 2012). In the control condition, we noticed that the density of MTs in the oocyte decreases from stage 5 to 7 (Sup Fig. 4 A-B). Furthermore, in *Khc* RNAi-mediated depletion, the MT density is slightly but significantly smaller than the control at stages 6B and 7. However, no significant difference could be detected between the control condition and the *Klc* RNAi-mediated depletion at any stage (Sup Fig. 4 A-B). In conclusion, since MT density is only weakly affected by the absence of Khc and not significantly by the absence of Klc, while both contexts affect nucleus positioning prior to migration, it is unlikely that the role of Kinesin-1 in this process is limited to its effect on MT stability.

**Figure 4.**
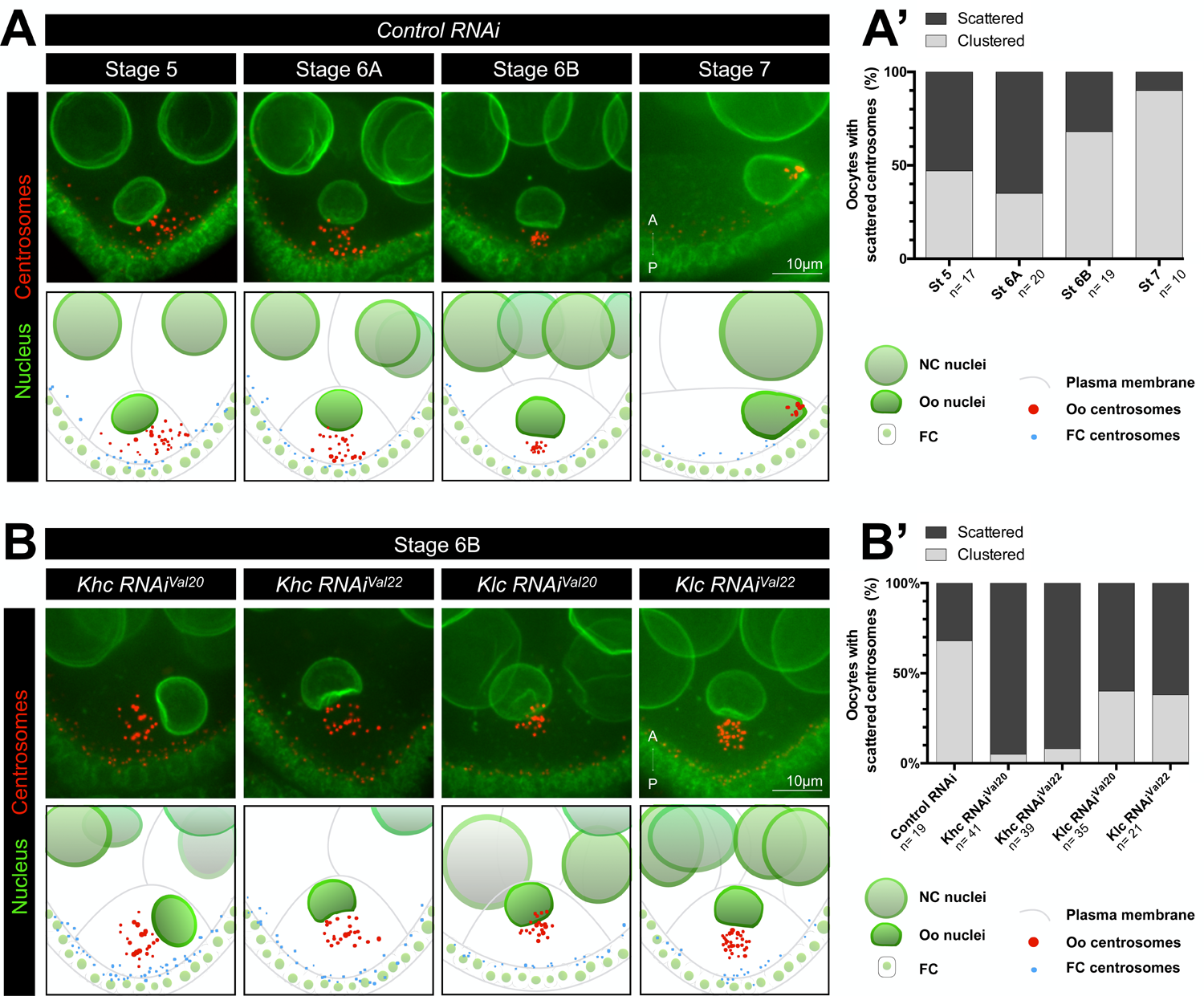
Khc and Klc are required for centrosome clustering at stage 6B. **(A-B)** (*Top row*) Representative Z-projection images of stage 5 to 7 egg chambers (A) and stage 6B (B), expressing *Fs(2)Ket-GFP* to label nuclei (green), *pubi-asl-tdTomato* to label centrosomes (red) and control RNAi (*UASp-CG12699 RNAi – See Methods*) (A) or the indicated RNAi (B) under the control of the *p_mat_αTub-Gal4* driver. The oocytes are oriented with anterior (A) at the top and posterior (P) at the bottom. Scale bar: 10µm. (*Bottom row*) Schematic diagrams of the image above, with oocyte centrosomes in red and follicular cell centrosomes in blue. **(A’-B’)** Quantification of oocytes categorized as scattered (black) and aggregate (gray) depending on centrosome distributions, at the different stages and for the different genotypes. n indicates the number of analyzed egg chambers. See Sup Table 1 for detailed values of the quantifications.

### Khc and Klc subunit are required for centrosomes clustering

Since we have previously reported two different and complementary cues involved in the migration of the *Drosophila* oocyte nucleus, i.e the MAP Mud and the centrosomes (Tissot et al., 2017), we next wondered whether Kinesin-1 affects either of these elements. When we quantified Mud asymmetry at the NE, we did not find any difference between the control condition and RNAi-mediated depletion of either *Khc* or *Klc* (Sup. Fig. 5). Then, we investigated a putative effect of Kinesin-1 on centrosomes. After oocyte specification, the centrosomes of the 15 NCs migrate through the ring canals into the oocyte, forming a cluster of at least 16 centrosomes (Bolívar et al., 2001; Nashchekin et al., 2021; Pimenta-Marques et al., 2016). Prior to nuclear migration, this cluster frequently coalesces in a compact structure in proximity of the nuclear side facing the oocyte posterior (Tissot et al., 2017; Zhao et al., 2012). We first monitored centrosome distribution by following the centrosomal protein Asterless (Asl) fused to RFP which allowed live imaging of the centrosomes in the developing oocyte (Tissot et al., 2017). Precise analysis in living conditions of control oocytes, revealed a switch in centrosome dispersion between stages 6A and 6B, when the nucleus centers itself in the oocyte. Whereas most centrosomes are scattered at stages 5 and 6A, most are clustered at stages 6B and 7 (Fig 4 A-A’). Live imaging experiments confirmed that the centrosomes, although in the vicinity of the nucleus, are dynamic and dispersed at early stages (Movies 1, 2). Then, while the nucleus centers itself in the oocyte, the centrosomes aggregate and remain clustered during migration (Movies 3, 4). We next investigated the requirement for Khc and Klc subunits in centrosome clustering in living conditions and found that both *Khc* and *Klc* RNAi-mediated depletions, impair centrosome clustering at stage 6B compared to control (Fig 4 B-B’ and Sup. Fig. 6). These results indicate that both subunits are required for centrosome clustering. Interestingly, a previous study has also reported this lack of clustering in absence of Klc (Hayashi et al., 2014). This Kinesin-1 effect upon centrosome clustering result is surprising, since plus-end directed MT-associated motors are involved in centrosome separation (Métivier et al., 2019), whereas centrosome clustering is usually ensured by minus-end directed MT-associated motors (Basto et al., 2008; Robinson et al., 1999). As Kinesin-1 often functions cooperatively with Dynein and the two motors are known to have interdependent functions (Duncan and Warrior, 2002; Januschke et al., 2002; Splinter et al., 2010), we then investigated if Kinesin-1 was required for Dynein localization at the centrosome. We found that, *Khc* RNAi-mediated depletion did not affect Dynein location at the centrosomes, as revealed by Dlic and Dynamitin, two subunits of the Dynein/Dynactin complex (Reck-Peterson et al., 2018) (Sup. Fig 7). This indicates that Kinesin-1 involvement in centrosome clustering is not simply due to the Dynein transport towards the centrosomes.

**Figure 5.**
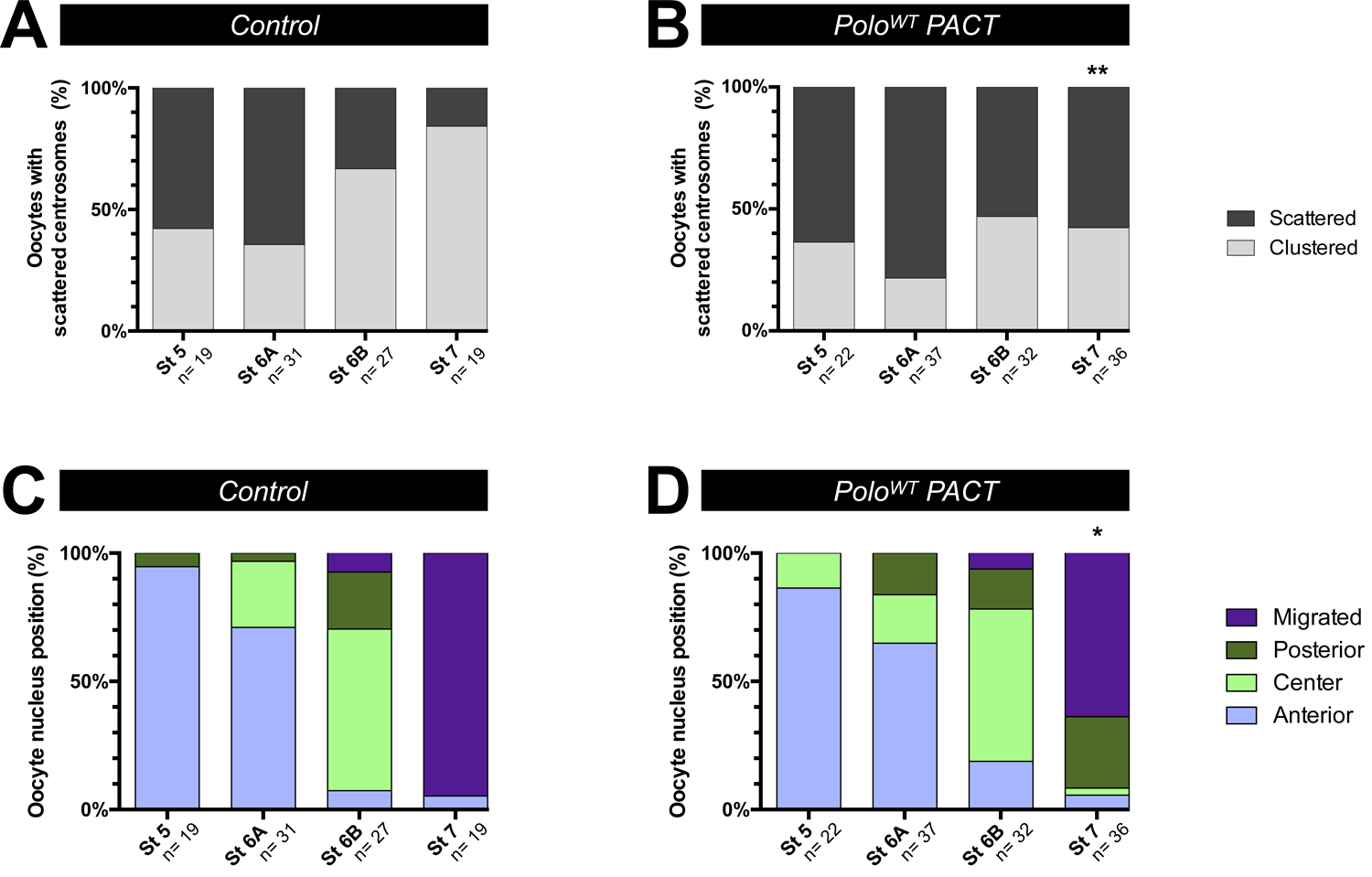
Preventing centrosome decay affects their clustering and nucleus positioning. **(A)** Quantification of oocytes categorized as scattered (black) and aggregate (gray) depending on centrosome distributions, at the different stages of control (*identical panel to* Figure 5A) and *UAS-Polo^WT^-PACT* over-expressing egg chambers under the control of the *p_mat_αTub-Gal4* driver. **(C-D)** Distribution of nucleus positions at the different stages has been quantified in the egg chambers quantified in A and B. Positions have been categorized and color-coded as anterior in pale blue, center in pale green, posterior in dark green and migrated in purple. n indicates the number of analyzed egg chambers. Chi2 test, *p < 0.05, **p = 0,005 compared to the control condition (per stage). See Sup Table 1 for detailed values of the quantifications.

**Figure 6.**
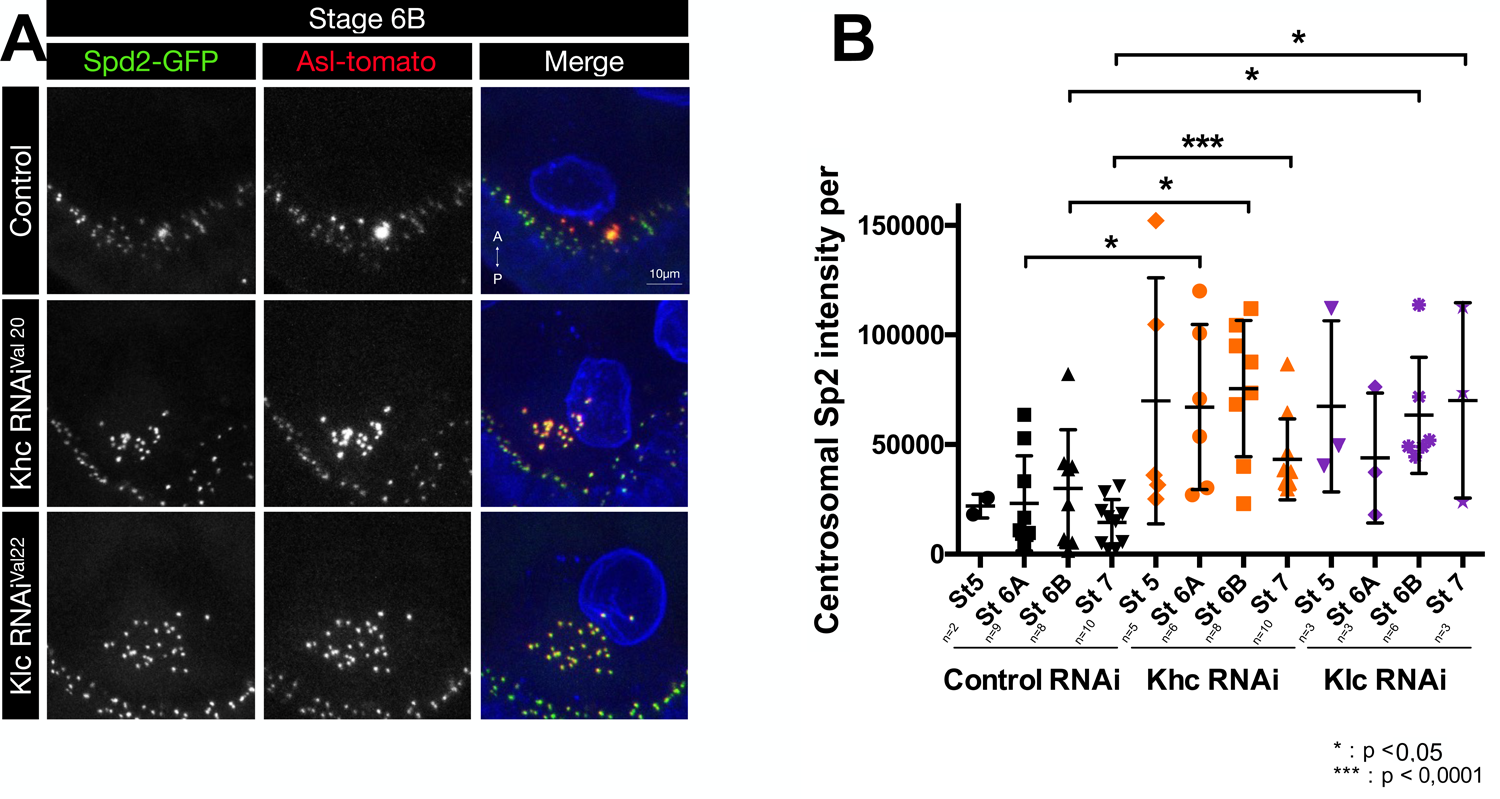
The level of SPD2 is increased at the centrosomes upon Kinesin-1 inactivation. **(A)** Representative Z-projection images of stage 6B egg chambers, expressing a control RNAi (*UASp-CG12699 RNAi*) or the indicated RNAi under the control of the *p_mat_αTub-Gal4* driver and *pubi-SPD2-GFP* (green) and *pubi-asl-tdTomato* to reveal centrosomes (red), and then stained with WGA to label nuclei. The oocytes are oriented with anterior (A) at the top and posterior (P) at the bottom. Scale bar: 10µm. **(B)** The quantification corresponds to the sum of intensity Spd2 at the centrosomes per oocyte. Mann-Whitney tests have been performed, and the significance is indicated (*: p<0,05 **: p<0,0001).

**Figure 7.**
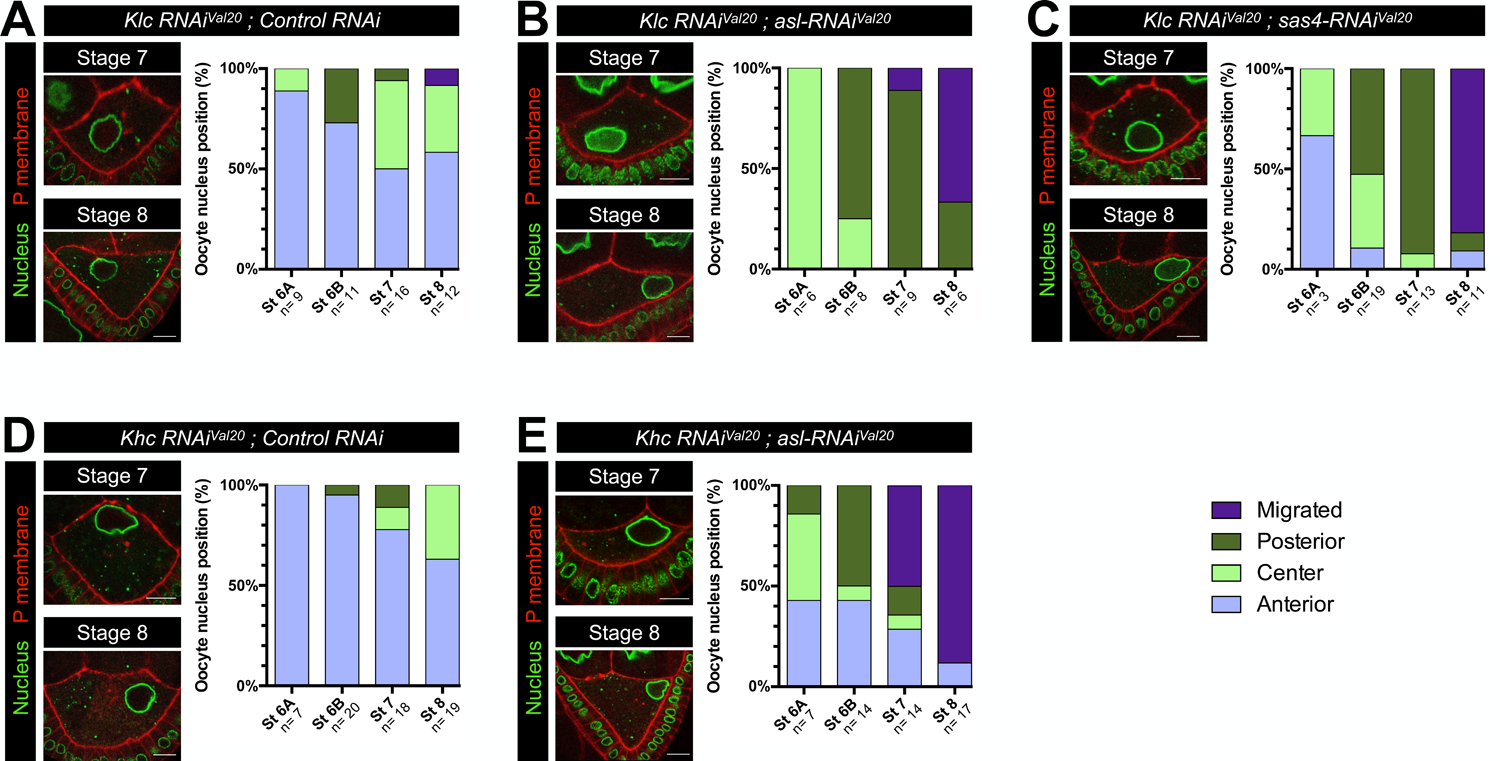
Centrosome inhibition and Kinesin-1 inactivation restore nuclear migration. Representative image of stage 7 and 8 egg-chambers and distribution of nucleus positions at the different stages. Positions have been categorized and color-coded as anterior in pale blue, center in pale green, posterior in dark green and migrated in purple. n indicates the number of analyzed egg chambers. **(A-E)** Expression of *Klc-RNAi^Val20^* (A-C) in combination with control RNAi (*UASp-Ap-RNAi – See Methods*) (A) *UASp-asl-RNAi^Val20^* (B) and *UASp-sas4-RNAi^Val20^* (C) or *Khc-RNAi^Val20^* in combination with control RNAi (*UASp-Ap-RNAi – See Methods*) (D) *UASp-asl-RNAi^Val20^* (E) using the *p_mat_αtub-Gal4* driver combined with *Fs(2)Ket-GFP* to label nuclei (green) and *pubi-PH^PLC∂1^-RFP* to label plasma membrane (red). See Sup Table 1 for detailed values of the quantifications.

### Kinesin-1 mediated centrosome clustering is needed for nuclear migration

Since centrosome clustering and nuclear centering occur concomitantly and both processes are affected by the loss of function of Kinesin-1, we wondered whether the two processes are related. Previous studies have reported that during the stages 5 to 7, all centrosomes, scattered or aggregated, are surrounded by pericentriolar material (PCM) and are therefore considered active i.e. they can organize microtubules (Januschke et al., 2006; Pimenta-Marques et al., 2016; Tissot et al., 2017; Zhao et al., 2012). Moreover, it has been shown that centrosomes exert pushing forces on the nucleus (Tissot et al., 2017; Zhao et al., 2012), therefore we reasoned that the anterior position of the nucleus results from centrosome generated forces applied to NE and centrosome clustering could be a sign of a decreasing centrosomal activity. Previously, it has been reported that centrosome elimination is a progressive process that starts at stage 6-7 with a decrease in PCM – centrosome association (Pimenta-Marques et al., 2016). Furthermore, it has been demonstrated that PCM decline is the consequence of Polo-like kinase 1 (Polo) decay from the centrosome. Hence, the ectopic tethering of an active form of Polo to centrioles, with a Pericentrin – AKAP450 Centrosomal Targeting (PACT) domain here after named Polo-PACT, is sufficient to prevent PCM loss and maintain active centrosomes (Pimenta-Marques et al., 2016).

To test our hypothesis that centrosome clustering observed in stage 6B could be a result of decreased centrosomal activity, we quantified centrosome clustering in oocyte expressing Polo-PACT transgene. Accordingly to our hypothesis, we found that Polo-PACT expression reduces the level of centrosome clustering, particularly at stage 7 (Fig 5 A, B), similarly to the defects observed when either *Klc* or *Khc* is inactivated by RNAi at the stages 6B and 7 (Fig 4 B, B’). To determine whether nucleus position defects observed in Kinesin-1 mutant backgrounds could be a consequence of defects in centrosome clustering, we assessed oocyte nucleus positioning when expressing Polo-PACT. In this context, we found that nuclear migration is significantly reduced compared to control at stage 7 (Fig. 5 C, D). This result highlights a link between the persistence of centrosome dispersion and a defect in nuclear migration. These results may suggest that a function of the Kinesin-1, with its two subunits Khc and Klc, is to promote centrosome clustering by promoting PCM removal. To directly test this possibility, we monitored SPD2, an essential PCM component and ASL levels at centrosomes (Fig. 6 A). By quantifying the intensity of SPD2-GFP signal using Imaris software (see methods), we found that at the stages 6A, 6B and 7 the level of SPD2 is significantly increased when *Khc* and *Klc* are inactivated (Fig. 6 B) compared to controls, showing that in absence of Kinesin 1 the level of PCM is increased at the centrosomes. This is further consistent with our hypothesis that clustered centrosomes have a reduced activity that is necessary for proper nuclear migration.

### Centrosome suppression and Kinesin-1 inactivation restore nuclear migration

Since nuclear migration defects in *Kinesin-1* inactivation context may be a consequence to excessive centrosome activity, we next asked whether Kinesin-1 inactivation associated with centrosome inactivation could restore the migratory capacity of the nucleus. In order to test this possibility, we induced centrosome inactivation through RNAi-mediated depletion of *Asl* (Sup. Fig 8) or *Sas-4*, two essential components for centrosome biogenesis (Blachon et al., 2008; Stevens et al., 2007), and analyzed the effect in combination with the inactivation of *Klc*. In *Klc-RNAi; asl-RNAi* double knockdown, as well as, *Klc-RNAi; sas4-RNAi* double knockdown, the positioning of the nucleus prior to its migration is shifted to the center and the posterior in comparison to the *Klc-RNAi* knockdown. In addition, the nuclear migration, although delayed, is significantly rescued at stage 8 (Fig. 7 A-C). Similarly, in *Khc-RNAi; asl-RNAi* double knockdown, the positioning of the nucleus at stage 6 is shifted to the center and the posterior compared to *Khc-RNAi* knockdown. Nuclear migration is also significantly rescued at stage 7 (Fig. 7 D-E). Altogether, these results indicate that centrosome inactivation restores the ability of the nucleus to center and to migrate when Kinesin-1 is inactivated. This further indicates, that Kinesin-1, with the involvement of its two subunits Klc and Khc, is essential for nuclear migration onset within *Drosophila* oocyte by controlling the level of centrosome activity and clustering.

## Discussion

In *Drosophila* oocyte, the MTs exert pushing forces required for nuclear migration (Tissot et al., 2017; Zhao et al., 2012). Previously, it was proposed that Kinesin-1 is not required for the nuclear migration in the oocyte but only for its maintenance in an asymmetrical position (Duncan and Warrior, 2002; Januschke et al., 2002; Palacios and St Johnston, 2002). In this study, we clearly show that Kinesin-1 is required for nuclear migration. Previous studies that reported that Khc and Klc are dispensable for nuclear migration did so on the basis of ovoD/FRT-mediated clonal analysis that did not show any positional defect of the nucleus at stages 6/7. We believe that it is possible that the ovoD dominant effect triggering oocyte degeneration (Chou and Perrimon, 1996), is not fully penetrant at stage 6/7 and thus the potential involvement of Khc and Klc subunits for nuclear migration has been overlooked (Duncan and Warrior, 2002; Januschke et al., 2002; Palacios and St Johnston, 2002). Importantly, with our RNAi-mediated analysis, as well as our GFP/FRT-mediated clonal analysis we have similar results to those previously published at stage 9 and beyond (Sup. Fig. 2).

Our results indicate that prior to its migration the nucleus moves from an anterior position to the center of the oocyte between the stages 5 and 6B. This suggests that, to migrate, the nucleus has to be centered in the oocyte. In addition, we report that Kinesin-1 controls this nuclear displacement, at least in part, by promoting centrosome clustering. This effect was not expected based on existing literature, as Kinesins and plus-end directed motors are generally involved in centrosome separation (Métivier et al., 2019) and instead minus-end directed motors bring the centrosomes closer (Robinson et al., 1999). We ruled out the possibility of an indirect effect on Dynein location, therefore a direct role of Kinesin-1 could be considered. In this regard, we also observed that in the *Drosophila* oocyte, increasing centrosome activity by over-expression of Polo kinase led to an impairment of centrosome clustering, hence suggesting that Kinesin-1 could have a role in PCM removal. Indeed, it has been previously shown that the decrease in Polo at the centrosome is responsible for the loss of PCM and the subsequent decrease in centrosome activity and disappearance (Pimenta-Marques et al., 2016). Accordingly, Kinesin-1 could either transport Polo or some PCM components away from active centrosomes. In this regard and interestingly, a recent work has reported that in *Drosophila* neuroblasts and squamous epithelial cells, Khc interacts directly with the PCM organizer Pericentrin-like protein (Plp) but also that Kinesin-1 is required for a differential distribution of Polo between the two centrosomes of a mitotic spindle (Hannaford et al., 2022). An attractive model would be that the PCM removed by Kinesin at the centrosomes is recycled to the previously described non-centrosomal MT sources in the oocyte, i.e the nucleus (Tissot et al., 2017) and the anterior cortex (Nashchekin et al., 2016). It is also interesting to note that the link between Kinesin-1 and centrosome activity has been previously suggested in the early *Drosophila* embryo where Kinesin-1 reduces centrosome motility (Winkler et al., 2015) and in the *C. elegans* zygote where Kinesin-1 prevents premature centrosome maturation (McNally et al., 2012).

In addition, our data revealed different requirements for the two Kinesin-1 subunits, Khc and Klc. While both proteins are required for nuclear centering and centrosome clustering, the nuclear positioning defect observed in the absence of Klc, but not Khc, is rescued at stage 9. This indicates that Khc independently of Klc can fulfill an additional function for nuclear positioning. It is quite striking that the delay for nuclear migration does not gradually appear after stage 7, as we would expect from a simple slowing down of the process, but is sharply occurring at stage 9. We and others have previously shown that the MT network in the oocyte undergoes dramatic rearrangement at stage 9 to organize bundles at the cell cortex (Drechsler et al., 2020; Januschke et al., 2006; Lu et al., 2016; Nashchekin et al., 2016; Parton et al., 2011; Trong et al., 2015). At this stage, the MTs generate a cytoplasmic flow which sustains an advection-based transport corresponding to an active transport induced by fluid flow (Drechsler et al., 2017; Drechsler et al., 2020; Ganguly et al., 2012; Loiseau et al., 2010; Williams et al., 2014). This process requires Khc activity, but not Klc, as in absence of Klc the cytoplasmic streaming is unaffected (Loiseau et al., 2010; Palacios and St Johnston, 2002; Williams et al., 2014). More recently, it has been identified that the origin of the cytoplasmic flow is generated by sliding of MT bundles against each other. Khc triggers the sliding by binding one MT with a C-terminal MT-binding site while walking along a second MT using its motor domain (Lu et al., 2016). This hypothesis does not exclude the alternative possibility that Khc has a different partner to perform its roles at a later stage, including the maintenance of the asymmetric position of the nucleus. Notably it was observed that the nucleus of egg chambers mutant for *ensconsin*, was not maintained in an asymmetric position (Metzger et al., 2012; Sung et al., 2008).

Previous works have highlighted the role of centrosomes as a cue to sustain the nuclear migration in the oocyte (Tissot et al., 2017; Zhao et al., 2012), even if centrosome inhibition does not prevent it (Stevens et al., 2007). One hypothesis is that very active and scattered centrosomes do not allow the oocyte nuclear migration, meaning that centrosomes would negatively regulate the nucleus migration. The seemingly contradiction of this results may simply underline the necessity of fine-tuning centrosome activity for nucleus migration. One possibility is that scattered centrosomes may exert uncoordinated forces on the nucleus that results to prevent nucleus migration. Therefore, a decrease in their activity would allow the centrosomes to cluster at the posterior of the NE and participate to the MT associated pushing forces required for nuclear migration.

## Materials and Methods

### Drosophila stocks and culture conditions

Drosophila stocks and crosses were maintained under standard conditions at 25°C. The following fly strains were used: *w^1118^* (Bloomington Drosophila Stock Center, BL#3605), *CantonS*, *p_mat_αtub-Gal4* BL#7062, *Fs(2)Ket-GFP* (Villányi et al., 2008), *pubi-PH-RFP* (Claret et al., 2014), *pubi-asl td-Tomato* gift of T. Avidor-Reiss (Chen and Yamashita, 2020; Gopalakrishnan et al., 2011), *p_mat_αTub-Khc-GFP* (Sung et al., 2008), *Klc-GFP* (Fly TransgenOme collection) (Sarov et al., 2016), *UASp-polo^WT^-PACT* (Pimenta-Marques et al., 2016), *pubi-DLic-GFP* (Baumbach et al., 2015), *p_mat_αTub -GFP-Dynamitin* (Januschke et al., 2002), *hsp-flp; FRT 42B pubi-GFP*, *hsp-flp; FRT 79D ubi-GFP* (gift from JR Huynh), *FRT 42B Khc^27^* (Januschke et al., 2002), *FRT 79D Klc^8ex94^* (Gindhart et al., 1998), *FRT 79D Klc^Saturn^*(Hayashi et al., 2014), *Jupiter-GFP* (Baffet et al., 2012), *pubi-spd2-GFP* (Conduit et al., 2014) Various informations needed for this study was found in FlyBase (Gramates et al., 2022).

### RNAi crosses

The following fly lines, generated from the TRIP project (Perkins et al., 2015) and obtained from the BDSC, have been used in this study: *UASp-Khc RNAi^Val20^* (attP2, Valium 20, HMS01519) BDSC #35770*, UASp-Khc RNAi^Val22^* (attP2, Valium 22, GL00330) BDSC #35409, *UASp-Klc RNAi^Val20-attP2^* (attP2, Valium 20, HMS00883) BDSC #33934*, UASp Klc RNAi^Val22^* (attP40, Valium 22, GL00535) BDSC #36795, *UASp-Klc RNAi^Val20-attP40^* (attP40, Valium 20, HMS02650) BDSC #42957*, UASp-ap RNAi^Val20^* (attP2, Valium 20, HMS02207) BDSC #41673*, UASp-Him RNAi^Val22^* (attP2, Valium 22, GL01183) BDSC #42809, *UASp-CG12699 RNAi^Val20^* (attP40, Valium20, HMS02833) BDSC#44111*, UASp-asl RNAi^Val20^* (attP2, Valium 20, HMS01453) BDSC #35039, *UASp-asl RNAi^Val22^* (attP40, Valium 22, GL00661) BDSC #38220, *UASp-sas4 RNAi^Val20^* (attP2, Valium 20, HMS01463) BDSC#35049 RNAi crosses were all performed using females from RNAi lines and maintained under standard conditions at 25°C. As control RNAi, we used lines expressing RNAi directed against genes not expressed in ovary, i.e *CG12699*, *apterous* (ap) and *Holes in Muscle* (Him) (Brown et al., 2014; Parisi et al., 2004). In addition, the lines were selected and used regarding the Valium plasmid used as well as the insertion point in the genome (attP2 or attP40).

### Heat-Shock Treatment for clonal analysis

Heat-shocks were carried out for 1 hr in a water bath at 37°C, 3 days in a row from L1 larvae.

### Immunostaining of the fly ovaries

1day old females were collected and put with fresh media for 30-48 hours at 25°C prior to dissection. Ovaries were dissected, fixed in PBS with 4% paraformaldehyde and incubated overnight at 4°C with primary antibodies in PBS with 0,1% Tween. Primary antibodies include: mouse α-Cut (DSHB #2B10, supernatant) at 1:50; mouse α-Hindsight (DSHB #1G9, supernatant) at 1:100; rabbit α-Mud (Izumi et al., 2006) at 1:1000; rabbit Anti-Khc (AKIN02-A, Cytoskeleton) 1:250. Ovaries were then incubated with secondary antibodies at room temperature for 2 hours. Secondary antibodies include: chicken α-mouse Alexa647 (Invitrogen, #A21463) at 1:100 and goat α-rabbit Alexa647 (Jackson Immunoresearch #111-605-144) at 1:200. For egg chambers that did not express a plasma membrane marker nor a nucleus marker, ovaries were respectively incubated with SiR-actin (Cytoskeleton TM) at 1:150 and with Wheat Germ Agglutinin (WGA) (Molecular Probes) at 1:200, at 4°C overnight. After washes, ovaries were mounted in Citifluor^TM^ (EMS). Images were captured using Zen software on a Zeiss 710 confocal microscope (488, 561, 640 nm lasers and the x40 or x63 objectives).

### Live imaging of egg chambers

To assess nucleus position, 1day old females were collected and put with fresh media for 30-48 hours at 25°C prior to dissection. Ovaries were dissected in Schneider medium and directly mount in halocarbon oil (Voltalef 10S) on a coverslip.

For egg chambers that did not express a plasma membrane marker, ovaries were first incubated with live-cell CellMask Deep Red (Invitrogen, #C10046) at 1:1000 diluted in Schneider medium room temperature for 10min.

To image the centrosome clustering, time-lapses of 4 hours (acquisition intervals of 5min) on living egg chambers have been performed, following previously described protocol (Loh et al., 2021).

Images were captured using Metamorph software on a Zeiss Axio Observer Z1 confocal microscope coupled with a spinning disk module CSU-X1 and a sCMOS camera PRIME 95 (488, 561, 640 nm lasers and a x63 oil immersion objective). In order to image the whole nucleus, 21 stacks along the z axis of a 1µm range were taken for each egg chamber.

### Image analysis, quantification and statistical analysis

All images were processed using the software Fiji (Schindelin et al., 2012).

#### Egg chamber staging

To stage the egg chambers, we measured nucleus diameters of the two closest nurse cells of the oocyte, using the z-section corresponding to the larger diameter of the considered nurse cells. These diameters were measured using the « Straight, segmented lines » tool on Fiji. In case the two nurse cells show diameters that can be categorized in different stages, we then took in account the shape and the size of the oocyte and the egg chamber aspect ratio as suggested by (Chen et al., 2019). Note that in the literature, over the years, there is a discrepancy about the stage at which the nucleus completes its migration, depending on the studies it could be either stage 7 or stage 8. In our case (Tissot et al., 2017), we had consistently named it stage 7 and our measures of stages 5, 6, 7 and 8 are in total accordance with what Bilder’s lab have called 6, 7, 8 and 9, respectively.

- Determination of nuclear position in the oocyte: Nucleus position was mainly assessed on a qualitative criterion, nevertheless in case of uncertainty, we have applied a quantitative method consisting in making the ratio of the distances: hemisphere anterior of the nucleus - anterior membrane (d*_ant_*) and hemisphere posterior of the nucleus - posterior membrane (d_post_); r = d*_ant_*/d*_post_*. If r < 0,5 the nucleus was categorized as “anterior”, if r > 2 the nucleus is determined as “posterior” and if 2 > r > 0,5, the nucleus is considered as “central”.

### Distribution of the centrosomes

To determine the distribution of the centrosome, we relied our qualitative analysis on two parameters: the number and the general spatial spreading in the oocyte. Centrosomes were considered as dispersed when either 10 centrosomes at least were clearly distinct or were spread over more than 5 µm.

### Measure of MT density

The MT signal intensity of the entire oocyte was measured on a Sum slices -projection and the contour of the oocyte was delimited with the tool « Freehand selections ».

### Signal analysis and quantification of SPD2 at the centrosomes in the oocyte

To determine the level of SPD2 at the centrosomes, we analyzed egg chambers expressing SPD2-GFP, Asl-td-tomato to reveal the centrosomes and WGA to highlight the nuclear envelope. Imaris software (Oxford Instruments) was used to segment a 5 μn diameter sphere at the level of the centrosome in which SPD2-GFP intensity was recorded. The quantification corresponds to the sum of centrosomal Spd2 intensity per oocyte. Mann-Whitney tests have been performed, and the significance is indicated (*: p<0,05 **: p<0,0001).

### Statistics

Bar plots as well as statistical tests were carried out with GraphPad Prism6.

## Acknowledgements

We would like to thank Déborah Dauvet, Frida Sanchez-Garrido and Kahina Sadaouli for the preliminary functional analyses of Khc and Klc in nuclear migration. We thank the ImagoSeine core facility of the Institut Jacques Monod, member of IBiSA and the France-BioImaging (ANR-10-INBS-04) infrastructure at the Institut Jacques Monod for their help and support. Monica Bettencourt-Dias, David Ish-Horowicz, Rippei Hayashi, Jean-René Huynh, Isabelle Palacios, FlyBase release (e.g. FB2022_03) used for data obtained and analysed using FlyBase, the Bloomington Drosophila Stock Center, Vienna Drosophila Research Center and the Developmental Studies Hybridoma Bank for fly stocks and reagents. We are grateful to Véronique Brodu, Paul Conduit, Nathaniel Henneman, Jean-Antoine Lepesant, Lionel Pintard for critical comments on the manuscript. We are grateful to the laboratory members for helpful discussions.

## Competing interests

The authors declare no competing or financial interests.

## Author contributions

The project was conceived by F. Bernard and A. Guichet. F. Bernard. and A. Guichet designed the experiments that were subsequently performed by M. Loh and F. Bernard. The data were analyzed by M. Loh, F. Bernard and A. Guichet. The project funding, administration, and supervision were provided by F. Bernard and A. Guichet. M. Loh prepared the figures. A. Guichet and F. Bernard wrote the manuscript, which was edited and reviewed by M. Loh.

## Fundings

This work was funded by the Fondation ARC (PJA-20181208148) pour la Recherche sur le Cancer, the Ligue Contre le Cancer Comité de Paris (RS20/75-17), the Association des Entreprises contre le Cancer (Grant Gefluc 2020 #221366) and by an Emergence grant from IdEx Université de Paris (ANR-18-IDEX-0001). M.L was supported by a fellowship from “Ministère de l’Education Nationale, de la Recherche et de la Technologie” (MENRT) obtained from the BioSPC doctoral school. F.B is supported by Université de Paris. A.G is supported by the CNRS.

---

**Movie 1.** Time-lapse movie of developing egg chamber from stage 5 to 6A expressing *Fs(2)Ket-GFP* to label nuclei (green), *pubi-PH^PLC∂1^-RFP* to label plasma membrane (red) and *pubi-asl-tdTomato* to label centrosomes (red). The nucleus is anteriorly positioned in the oocyte and the centrosomes are scattered between the nucleus and the posterior membrane of the oocyte. Scale bar: 10µm. Time is indicated (h:min)

**Movie 2.** Crop of Movie 1 focusing on the oocyte region of the egg chamber. Scale bar: 10µm. Time is indicated (h:min).

**Movie 3.** Time-lapse movie of developing egg chamber from stage 6B to 7 expressing *Fs(2)Ket-GFP* to label nuclei (green), *pubi-PH^PLC∂1^-RFP* to label plasma membrane (red) and *pubi-asl-tdTomato* to label centrosomes (red). The centrosomes aggregate at the posterior of the centered nucleus, and follow the nucleus during its migration from the center of the oocyte to the cortex antero-lateral of the oocyte. Scale bar: 10µm. Time is indicated (h:min).

**Movie 4.** Crop of Movie 3 focusing on the oocyte region of the egg chamber. Scale bar: 10µm. Time is indicated (h:min).

## Notes

### Competing Interest Statement

The authors have declared no competing interest.

### Summary of Updates

To test the involvement of Kinesin-1 in regulating centrosomal activity, we aimed to assess the levels of PCM components in control RNAi and Kinesin RNAi. We quantified Spd2, an essential PCM component, by monitoring Spd2-GFP level at the oocyte centrosomes, revealed by Asl-tomato, in control RNAi, Khc RNAiVal20, and Klc RNAiVal22. These results are now presented in the manuscript as Figure 6. Thus, in absence of Kinesin-1, SPD2 an essential component of the pericentriolar material is increased at the centrosomes, suggesting that Kinesin-1 associated defects result from a failure to reduce centrosome activity.

