## Supplementary Figures for "Kinesin-1 promotes centrosome clustering and nuclear migration in the *Drosophila* oocyte"

**Supplementary Figure legends**

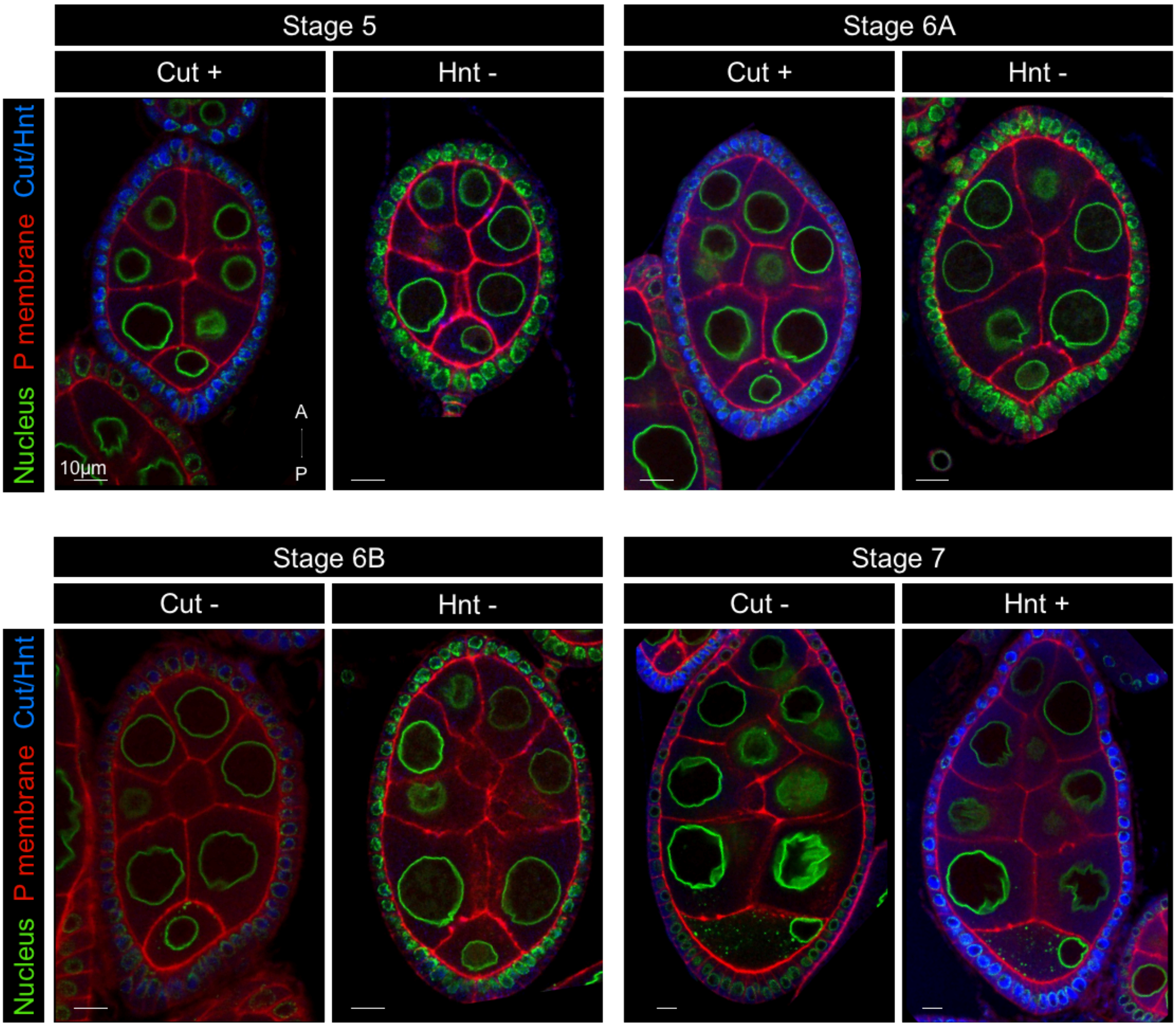


**Supplementary Figure 1. Expression of Cut and Hnt before and after the oocyte nuclear migration**

Stage 5 to 7 egg chambers, expressing *Fs(2)Ket-GFP* to label nuclei (green) and *pubi-PH*$^{PLC\partial 1}$-*RFP* to label plasma membrane (red), stained with Cut or Hindsight (Hnt) antibodies (blue). At stage 5 and 6A, the follicular cells show a clear signal for Cut but not for Hnt. At stage 6B, the follicular cells are mostly negative for Cut and Hnt. At stage 7, the follicular are negative for Cut and positive for Hnt staining. The egg chambers are oriented with anterior (A) at the top and posterior (P) at the bottom. Scale bar : 10μm.

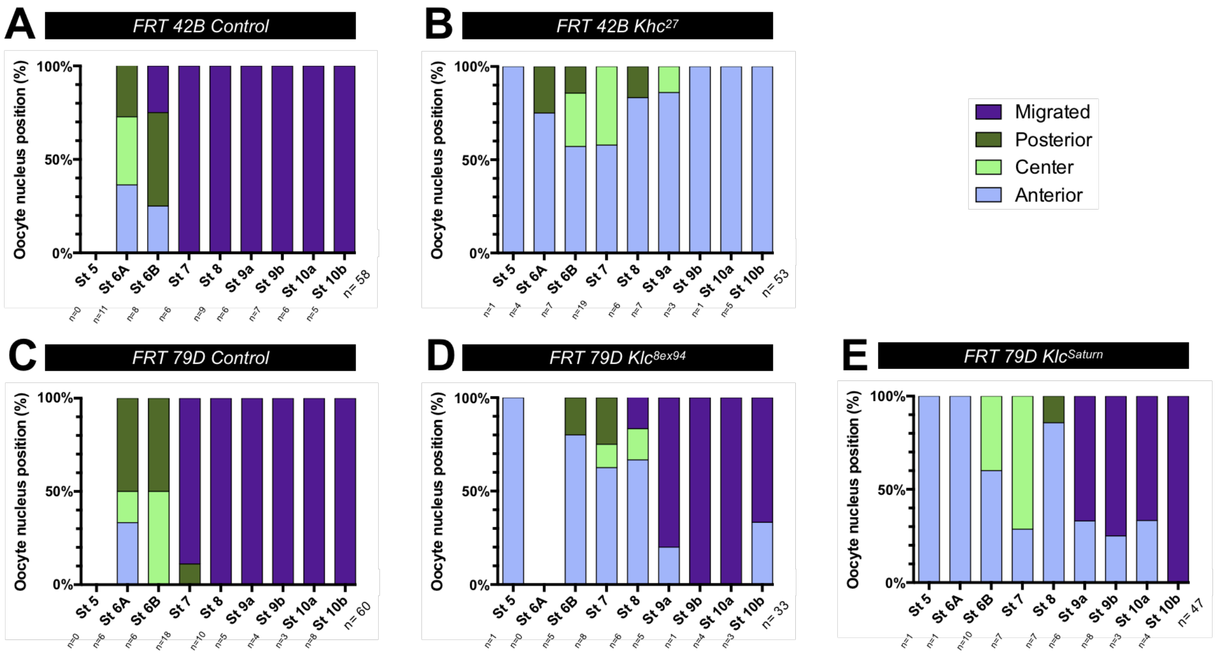


**Supplementary Figure 2. Khc and Klc are differentially required for nuclear migration. (A-E)** Distributions of nucleus positions at the different stages. Positions have been categorized and color- coded as anterior in pale blue, center in pale green, posterior in dark green and migrated in purple. n indicates the number of analyzed egg chambers. GFP/FRT clonal analysis of nucleus positions in control egg chambers $Khc^{27}$ heterozygous (A) and $Khc^{27}$ mutant (B), *Klc* heterozygous (C), $Klc^{8ex94}$ mutant (D) and $Klc^{Saturn}$ (E) mutant egg chambers. See sup table 1 for detailed values of the quantifications.

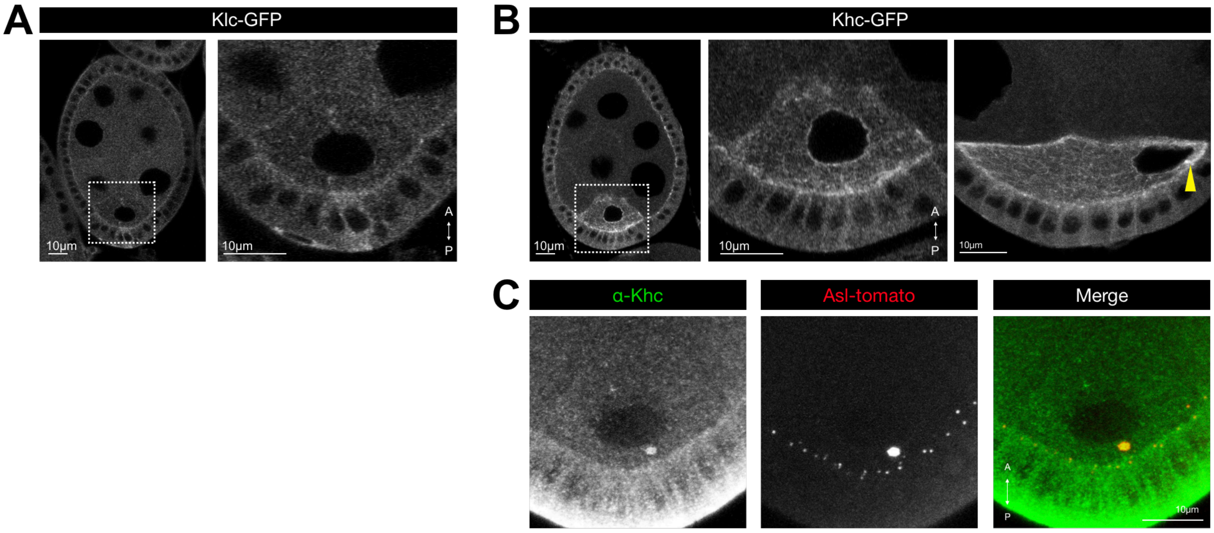


**Supplementary Figure 3. Khc and Klc localize differently in the oocyte.** **(A)** Stage 6B egg chamber expressing *ptub-Khc-GFP* in a *Khc*$^{27}$ homozygous background (left), and higher magnification corresponding the square display on A (right) (n=10) **(B)** Stage 6B of *Klc-GFP* egg chamber (left), and higher magnification corresponding the square display on A (right) (n=18). **(C)** Stage 6B egg chamber expressing *pubi-asl-tdTomato* to label centrosomes and stained with Khc antibodies (n=6).

Scale bars : 10μm.

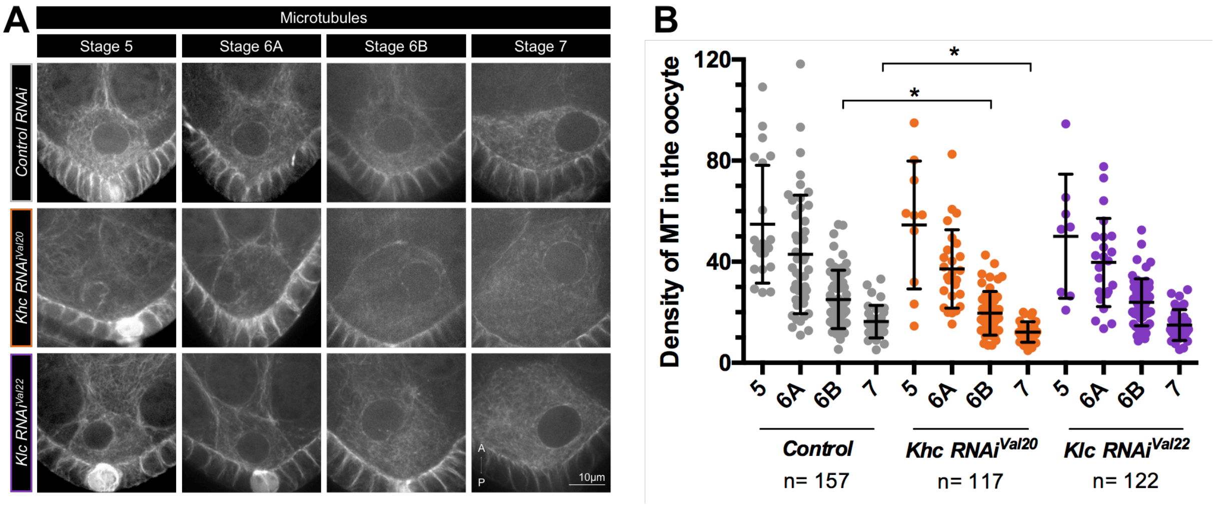


**Supplementary Figure 4. Khc and Klc impact differently the MTs.**

**(A)** Representative examples of oocytes of stage 5 to 7 egg chambers expressing *Jupiter-GFP,* to label the MTs, and control RNAi (*UASp-CG12699 RNAi – See Methods*) (top row), *Khc RNAi$^{Val20}$* (middle row) and *Klc RNAi$^{val22}$* (bottom row) under the control of the *$p_{mat}\alpha$tub-Gal4* driver. The oocytes are oriented with anterior (A) at the top and posterior (P) at the bottom. Scale bar : 10µm. **(B)** Quantification of MT density in oocytes of the indicated stages and genotypes. Mann-Whitney test, *$p < 0.05$. n indicates the number of analyzed egg chambers. See Sup Table 1 for detailed values of the quantifications.

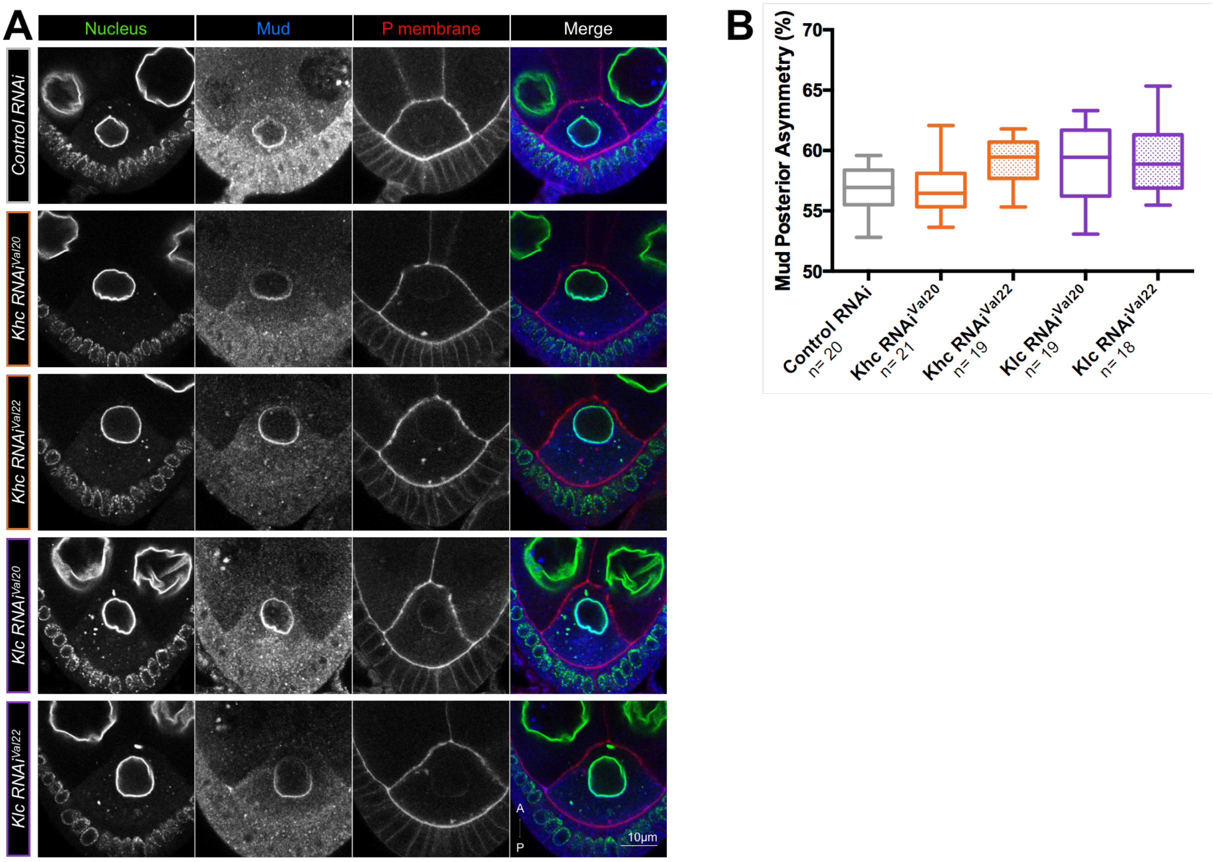


**Supplementary Figure 5. Kinesin-1 mutation does not affect Mud asymmetry at the nuclear envelope.**

**(A)** Representative images of stage 6B egg chambers, expressing *Fs(2)Ket-GFP* to label nuclei (green) and *pubi-PH*$^{PLC\delta1}$*-RFP* to label plasma membrane (red) and the indicated RNAi stained with Mud antibodies (blue). **(B)** Quantification of the percentage of signal intensity in the posterior hemispheres at the nuclear envelope of oocytes of the indicated genotype. A percentage higher than 50% corresponds to a posterior enrichment (*see Methods*). n indicates the number of analyzed egg chambers. Scale bar : 10µm. See sup table 1 for detailed values of the quantifications.

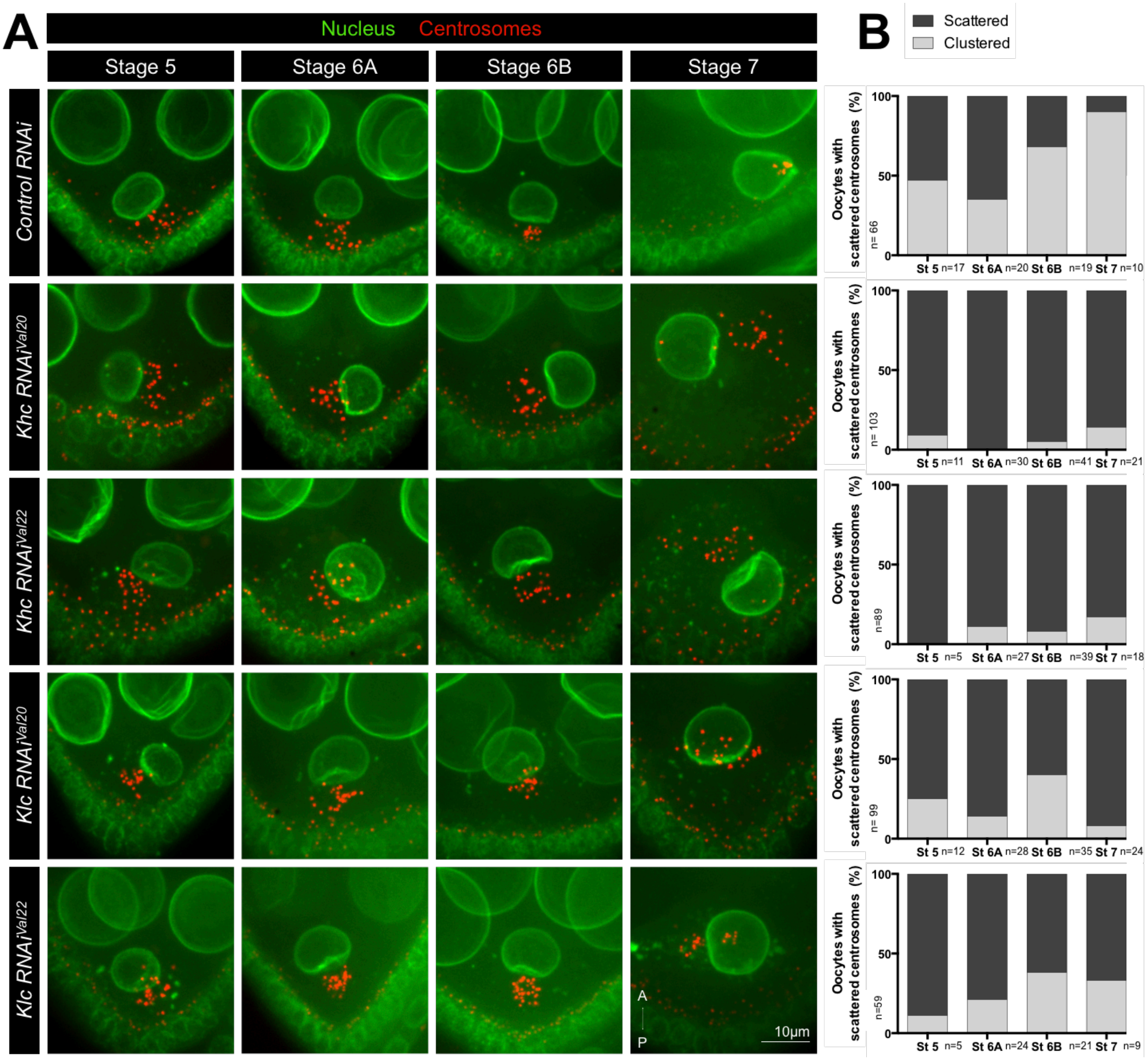


**Supplementary Figure 6. Clustering of the centrosomes is affected by Khc and Klc mutation.**

**(A)** Representative images of stage 5 to 7 egg chambers, expressing *Fs(2)Ket-GFP* to label nuclei (green), *pubi-asl-tdTomato* to label centrosomes (red) and the indicated RNAi under the control of the $p_{mat}\alpha Tub\text{-}Gal4$ driver. Scale bar : 10µm. **(B)** Quantification of oocytes categorized as scattered (black) and aggregate (gray) depending on centrosome distributions, at the different stages for each genotype. See sup table 1 for detailed values of the quantifications.

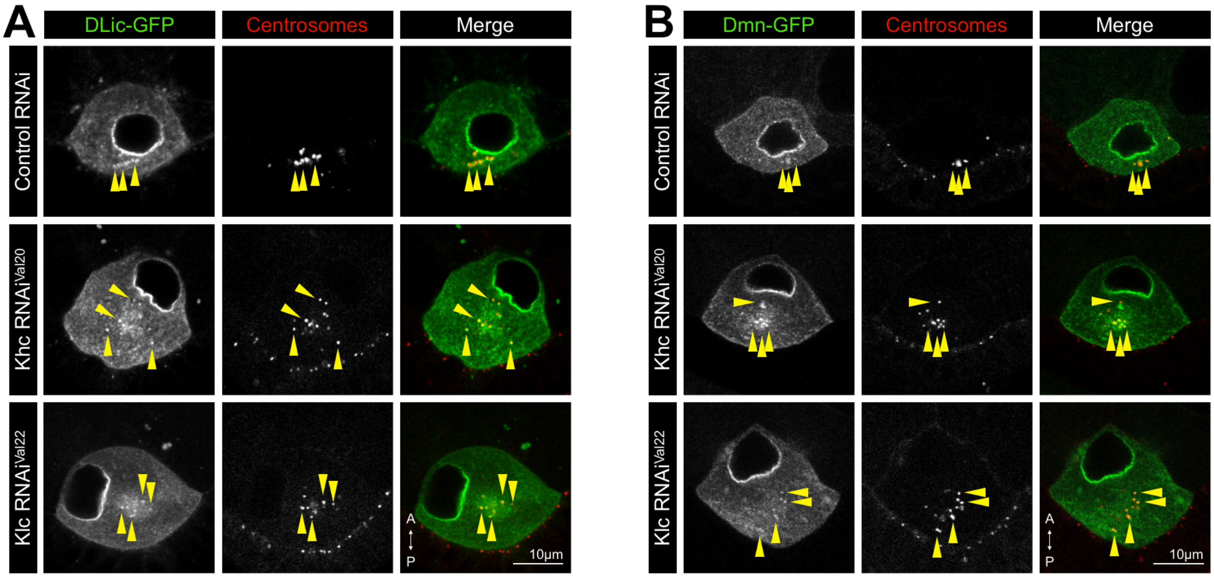


**Supplementary Figure 7. Kinesin-1 depletion does not affect Dynein motor localization from the centrosomes.**

**(A-B)** Oocytes from stage 6B egg chambers expressing *pubi-DLiC-GFP* (A) or $p_{mat}\alpha tub$ - *GFP-Dynamitin* (B, Dmn-GFP), *pubi-asl-tdTomato* to label centrosomes (red) and control RNAi (*UASp-Him RNAi – See Methods*) (top row) (DLic-GFP: n=36 ; Dmn-GFP: n=34), *Khc RNAi*$^{Val20}$ (middle row) (DLic-GFP: n=35 ; Dmn-GFP: n=31) and *Klc RNAi*$^{val22}$ (bottom row) (DLic-GFP: n=32 ; Dmn-GFP: n=35) under the control of the $p_{mat}\alpha Tub$-*Gal4* driver. The yellow arrowheads indicate the centrosomes. Scale bars : 10µm.

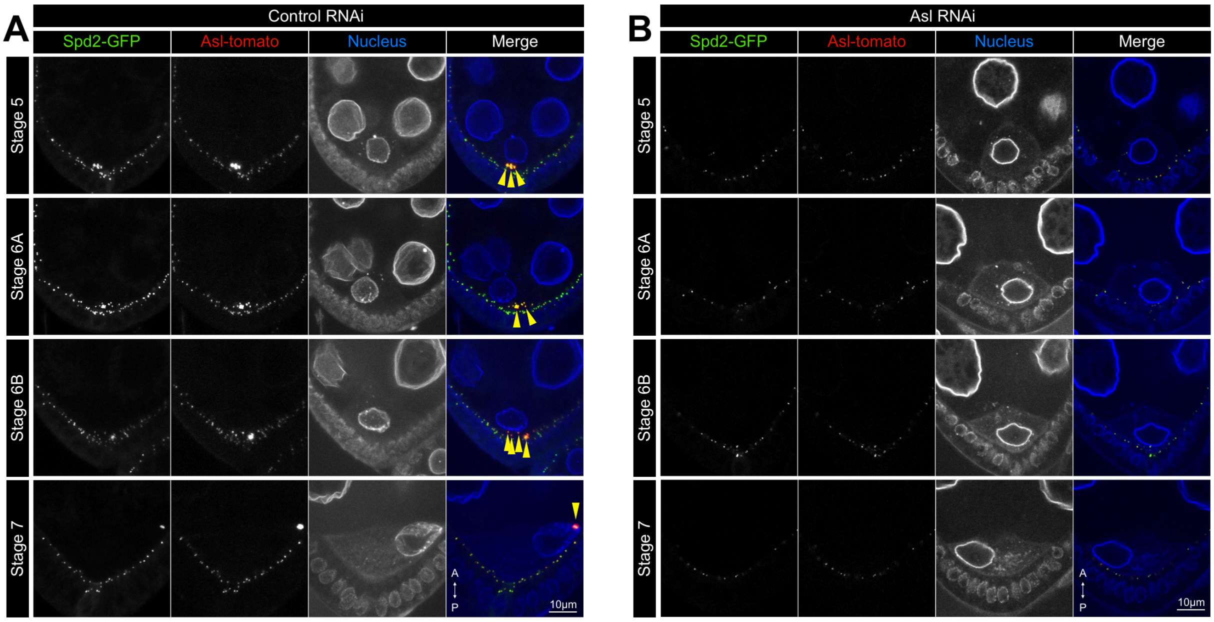


**Supplementary Figure 8. Asl depletion affects centrosome maintenance in the oocyte.** **(A)** Representative images of stage 5 to 7 egg chambers, expressing *pubi-SPD2-GFP* to label PCM (green), *pubi-asl-tdTomato* to label centrosomes (red) and control RNAi (*UASp-CG12699 RNAi*) under the control of the *mat-*α*Tub-Gal4* driver (n=40). **(B)** Representative images of stage 5 to 7 egg chambers, expressing *pubi-SPD2-GFP* to label PCM (green), *pubi-asl-tdTomato* to label centrosomes (red) and *asl-RNAi* $^{Val22}$ under the control of the $p_{mat}\alpha Tub$-*Gal4* driver (n=22). Scale bar: 10µm.
